# DTI versus NODDI White Matter Microstructural Biomarkers of Alzheimer’s Disease

**DOI:** 10.1101/2025.08.22.671829

**Authors:** Kenny Liou, Sophia I. Thomopoulos, Hannah Yoo, Yuhan Shuai, Sasha Chehrzadeh, Arvin Arani, Bret Borowski, Robert I. Reid, Clifford R. Jack, Michael W. Weiner, Neda Jahanshad, Paul M. Thompson, Talia M. Nir, Alzheimer’s Disease Neuroimaging Initiative

## Abstract

Diffusion MRI (dMRI) is a powerful tool to assess white matter (WM) microstructural abnormalities in Alzheimer’s disease (AD). The fourth phase of the Alzheimer’s Disease Neuroimaging Initiative (ADNI) now includes multiple multishell dMRI protocols, enabling both traditional and advanced dMRI model analyses. There is a need to evaluate whether multishell data offer deeper insights into WM pathology in AD than more widely available single-shell data by overcoming single-shell model limitations. Here, we fit single-shell DTI and multishell NODDI to dMRI data from 533 ADNI3/4 participants to assess their sensitivity to key clinical indicators of AD such as cognitive impairment, amyloid-beta and tau PET burden. Overall, we found that NODDI offered no major advantages in detecting cognitive impairment and tau pathology, but NODDI was marginally more sensitive to amyloid pathology.

## I. Introduction

Diffusion MRI (dMRI) is a powerful imaging technique for evaluating brain tissue, providing valuable insights into white matter (WM) microstructure. Its sensitivity to WM abnormalities has made it a key tool in the study of neurodegenerative diseases such as Alzheimer’s disease (AD) [1]. It is therefore critical to understand the utility of dMRI as a sensitive biomarker for key AD pathology, including amyloid-beta (Aβ) and tau, in addition to downstream cognitive impairment.

Historically, single-shell dMRI acquisitions have been the most used due to their generally shorter scan times and less demanding hardware requirements. This has led to the widespread use of diffusion tensor imaging (DTI) [2], a model that characterizes Gaussian diffusion from single-shell acquisitions. Numerous DTI studies have identified AD-related WM abnormalities, such as greater diffusivity and reduced anisotropy, in key brain regions like the corpus callosum, cingulum bundle, and fornix [3,4]. However, DTI has significant limitations. It cannot accurately model complex fiber architectures, such as crossing fibers, and its metrics often lack biological specificity, making it difficult to precisely link observed changes to underlying tissue properties [5]. Multishell dMRI protocols address these limitations by capturing both Gaussian and non-Gaussian diffusion signals, allowing finer characterization of WM. In particular, biophysical multishell models, such as neurite orientation dispersion and density imaging (NODDI) [6], can parse signal contributions from various tissue compartments including the intracellular compartment and free water. There is a need to evaluate whether multishell data offers deeper insights into WM pathology in AD by moving beyond the limitations of DTI.

The Alzheimer’s Disease Neuroimaging Initiative (ADNI) is an ongoing large-scale, longitudinal, multicenter study dedicated to identifying brain imaging, clinical, cognitive, and molecular biomarkers of AD and aging [7]. Now in its fourth phase, ADNI has expanded from a single multishell protocol to multiple multishell dMRI protocols across GE, Philips and Siemens scanners, enabling both traditional and advanced dMRI model analyses. In this study, we compared the sensitivity of ADNI DTI and NODDI WM measures to several key AD indicators: Aβ burden, tau pathology, and clinical measures of cognitive impairment. We also assessed whether sex moderates these effects. This is especially important given the significant sex differences in AD, where females not only have a higher prevalence than males, but also exhibit distinct pathological and cognitive profiles [8, 9]. We hypothesized that NODDI WM measures would provide better biological specificity and greater sensitivity to AD metrics and that larger effects would be detected in females than males.

## II. Methods

### A. Subject and Image Acquisition

Baseline 3 T T1-weighted (T1w) MRI and multishell dMRI data were downloaded from the ADNI database (https://ida.loni.usc.edu/). In total, we analyzed dMRI data from 533 participants across ADNI3/4: 341 cognitively normal elderly controls (CN), 149 mild cognitive impairment cases (MCI), and 43 dementia cases (**Table 1)**. All participants were scanned with one of four ADNI3/4 multishell dMRI protocols (**Table 2**). Key clinical indicators of AD, specifically Clinical Dementia Rating sum-of-boxes (CDR-sob; N=529) [10], Aβ-PET centiloids (CLs; N=213) [11], and tau-PET standardized value uptake ratios (SUVRs; N=186) [12] were obtained when available. Cortical Aβ-PET (18F-florbetaben, 18F-florbetapir, 18F-NAV4694) CLs were derived using methods described in [13]. Tau-PET (18F-flortaucipir) burden was defined using the medial temporal SUVR normalized using measures from the inferior cerebellar grey matter.

**Table 1.**
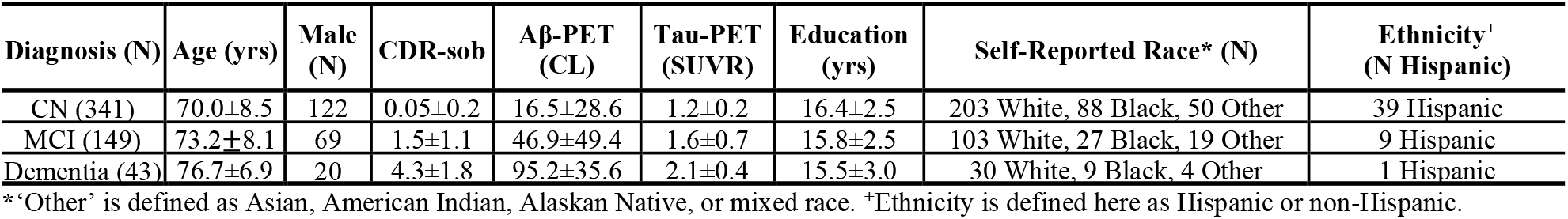
Subject Demographic and Clinical Information.

**Table 2.**
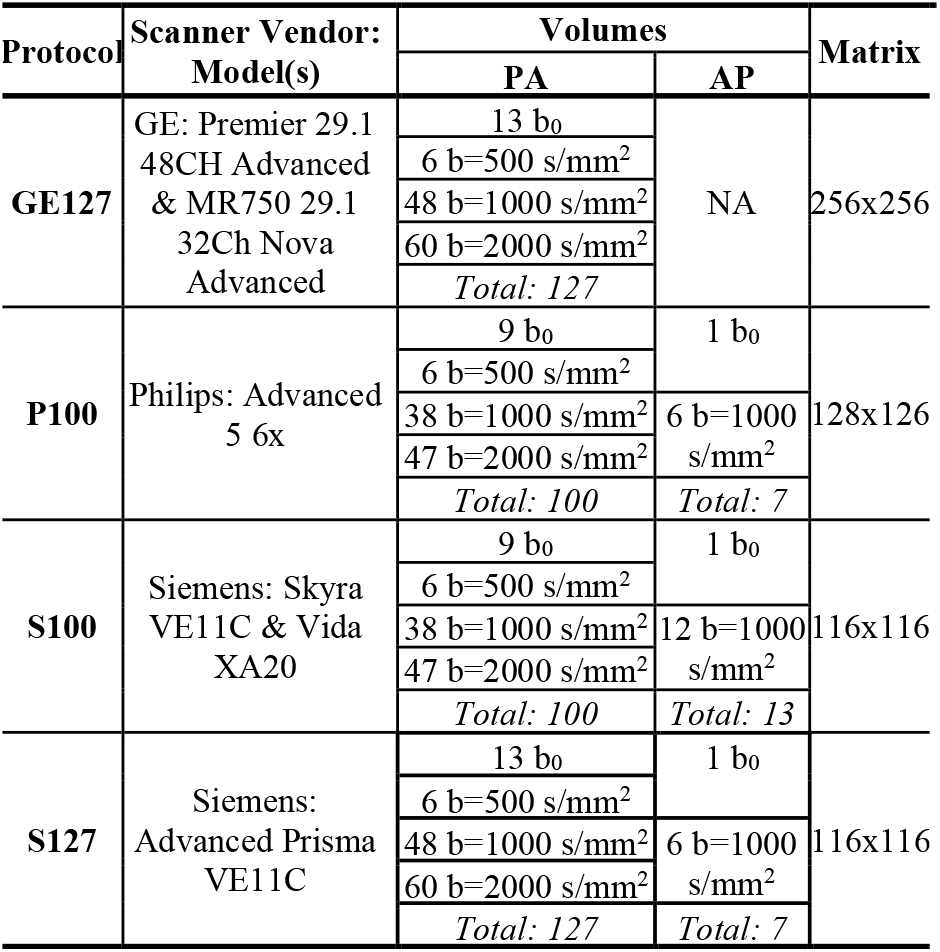
Available ADNI3 and ADNI4 multishell acquisition protocols. All dMRI were acquired with an isotropic 2×2×2 mm^3^ voxel size. GE127 dMRI were zero-padded in *k-*space during acquisition to a 0.9×0.9×2 mm^3^ voxel size.

### B. Image Preprocessing

ADNI3 dMRI were preprocessed as in [14]. As in ADNI3, all dMRI were denoised using principal component analysis (PCA) techniques [15,16] and corrected for Gibbs ringing [17]. For ADNI4 only, susceptibility distortions were corrected using FSL’s *topup* [18] using concatenated anterior-posterior (AP) and posterior-anterior (PA) scans. The concatenated AP-PA images were carried through the remaining preprocessing and analysis pipelines. GE multishell scans were excluded from *topup* since distortions were corrected in-scanner. Extra-cerebral tissue was removed, and eddy correction performed with FSL’s *eddy* tool, including outlier and slice timing corrections [19, 20, 21]. The resulting dMRI were corrected for B1 field inhomogeneities and warped to the subjects’ respective T1w (preprocessed with the standard FreeSurfer pipeline [22]) using ANTs; the registration was subsequently inverted to bring the T1w to the native DWI space [14]. All dMRI and T1w images were visually checked for quality assurance.

### C. DTI and NODDI Extraction

DTI was fit on the subset of b0 and b=1000 s/mm^2^ dMRI volumes and NODDI was fit across all dMRI volumes. DTI was computed using FSL’s weighted least squares function and NODDI was computed using DiPy [23]. Fractional anisotropy (FA), as well as mean (MD), axial (AxD), and radial diffusivity (RD) maps were derived from DTI. Intracellular volume fraction (ICVF), isotropic volume fraction (ISOVF), and orientation dispersion index (ODI) maps were extracted from NODDI. As in [14], the JHU ICBM-DTI-81 [24] atlas FA was warped to each subject’s FA. The transformations were applied to stereotaxic JHU WM atlas labels (JHU MNI single subject WMPM1) [24] using nearest neighbor interpolation. Mean DTI and NODDI measures were extracted from 31 WM regions of interest (ROIs), averaged across the left and right hemispheres (**Table 3**).

**Table 3.**
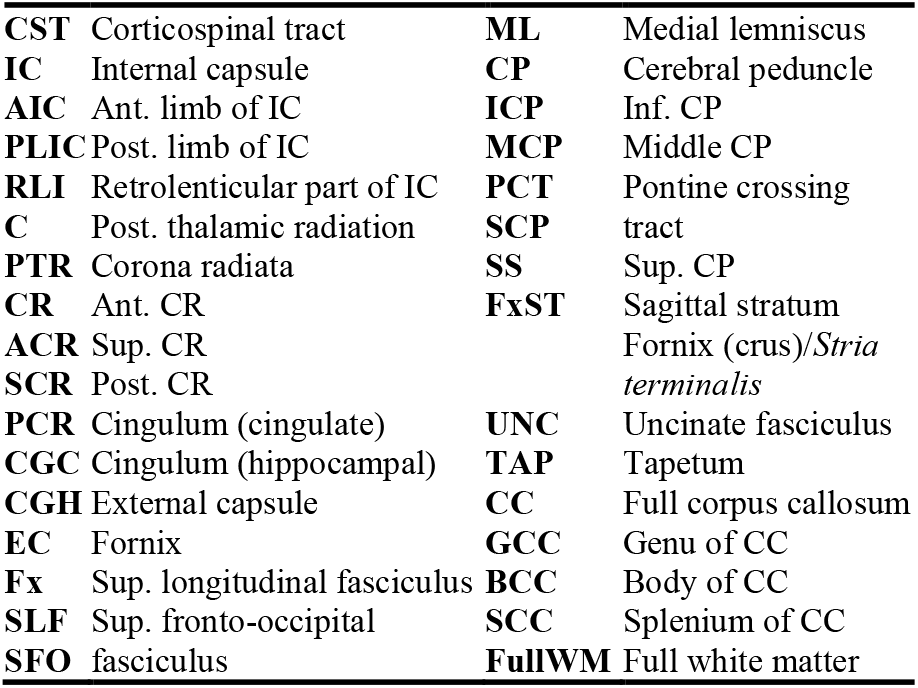
Index of 31 JHU atlas WM ROIs.

### D. Statistical Analyses

Linear mixed effects models were used to assess associations between regional dMRI measures and four key AD indicators: (1) CDR-sob, (2) an MCI versus CN diagnosis, Aβ-PET CLs, and (4) tau-PET SUVRs. We also tested whether sex moderated the effect between AD indicators and dMRI measures. Fixed effect covariates included age, sex, age-by-sex interactions, education, self-reported race, and ethnicity. We additionally covaried for diagnosis in Aβ and tau analyses. dMRI protocol and preprocessing scheme was modeled as a random effect. Effect sizes were estimated using partial *d* and partial correlation *r* for categorical and continuous AD metrics, respectively. The false discovery rate (FDR) procedure [25] was used to correct for multiple comparisons across 31 ROIs.

## III. Results

### A. DTI and NODDI Associations with AD Outcomes

Lower FA and ICVF, and higher AxD, MD, RD, and ISOVF were widely associated with greater cognitive impairment (i.e., higher CDR-sob and an MCI diagnosis; PFDR<0.05; **Fig. 1A, B**). While effects were detected in the Full WM, the strongest effects localized to the CC and the limbic tracts including the CGH, UNC, and Fx. Fewer associations were seen with ODI. Greater Aβ burden was linked to higher AxD, MD, ISOVF, and lower ODI across the IC (ISOVF), SCP (AxD, ODI), and Fx (ODI) (**Fig. 1C**). Tau burden was associated with higher AxD in the CC, FA and ICVF in the RLIC, and ISOVF in the CGC. Greater tau was also associated with lower RD in the RLIC and ODI in the CC (**Fig. 1D**).

**Figure 1.**
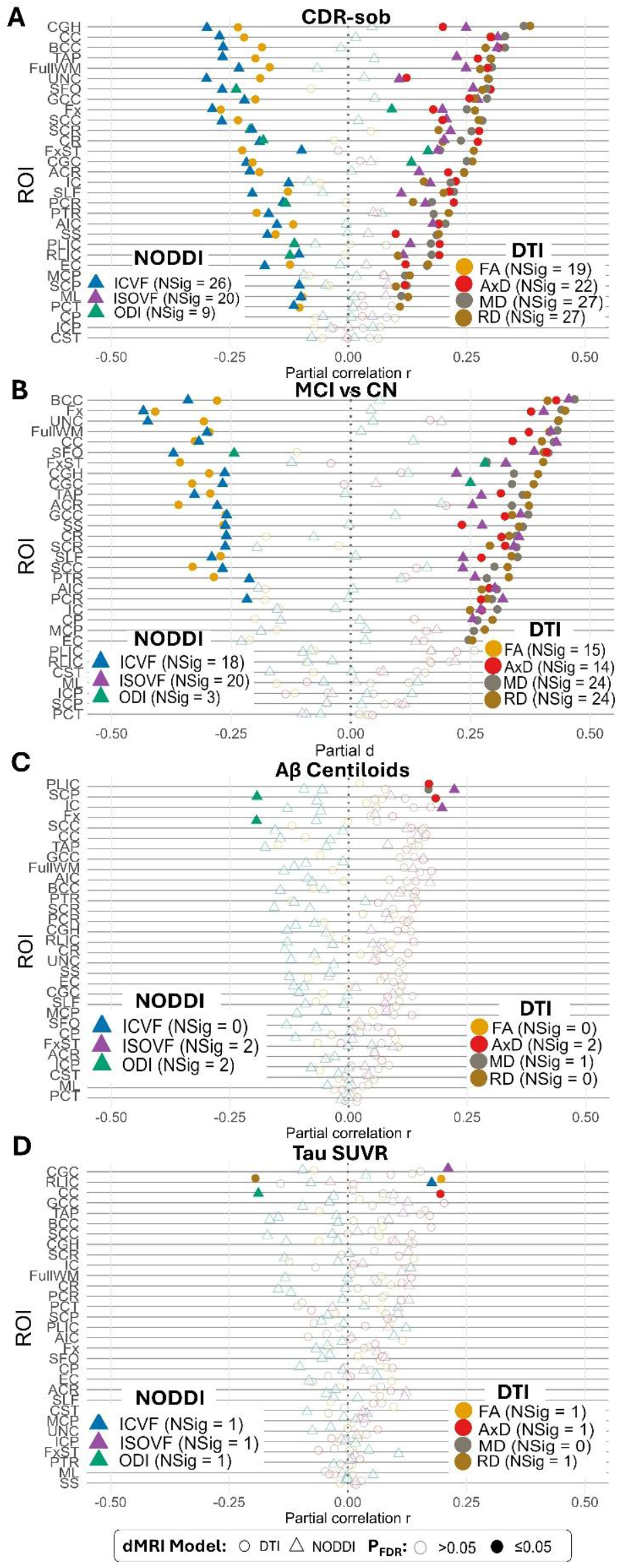
Effect sizes for associations between regional DTI and NODDI measures and (**A**) CDR-sob, (**B**) MCI diagnosis, (**C**) Aβ-PET CLs, and (**D**) tau-PET SUVRs. Significant associations that survive multiple comparisons corrections are opaque (*P*_FDR_<0.05).

### B. Sex-Specific Patterns in DTI and NODDI measures

Few sex-by-cognitive impairment interactions were found: no CDR-by-sex interactions were detected and only 3 dMRI measures showed significant MCI-by-sex interactions (**Table 3**). Males showed higher AxD and MD in the EC and higher AxD in the UNC compared to females. In contrast, a greater number of sex-by-PET pathology interactions were detected (**Table 3**). Again, with greater Aβ and tau load, overall males showed greater diffusivity and ISOVF, and lower FA, ICVF and ODI compared to females. As an example of sex-by-PET trends, scatter plots of FA in the cingulum illustrate that males showed a steeper decline in anisotropy compared to females with increasing Aβ (**Fig 2A**) and tau load (**Fig 2B**).

**Table 3.** Number of ROIs showing significant sex interactions per AD indicator and dMRI measure (*P*_FDR_<0.05).

| Test | Total | DTI |  |  |  | NODDI |  |  |
| --- | --- | --- | --- | --- | --- | --- | --- | --- |
|  |  | FA | MD | RD | AxD | ICVF | ISOVF | ODI |
| CDR-sob | 0 | 0 | 0 | 0 | 0 | 0 | 0 | 0 |
| CN vs MCI | 3 | 0 | 1 | 0 | 2 | 0 | 0 | 0 |
| Aβ-PET | 20 | 2 | 4 | 5 | 2 | 2 | 4 | 1 |
| Tau-PET | 23 | 6 | 4 | 5 | 1 | 1 | 4 | 2 |

**Figure 2.**
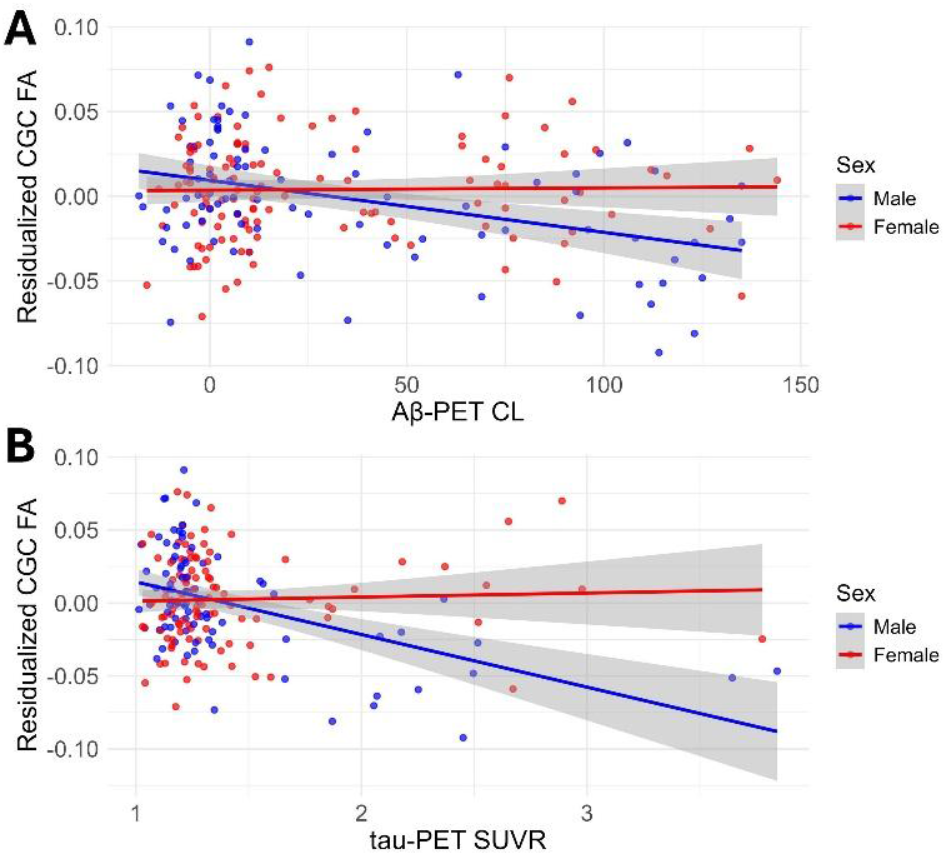
Cingulum (CGC) FA residuals illustrate that males showed a steeper decline in anisotropy compared to females with increasing (**A**) Aβ and (**B**) tau load.

## IV. Discussion

Overall, NODDI offered no major advantages in detecting measures of clinical impairment and tau pathology. The largest effects were often found in limbic and temporal lobe regions that are vulnerable in AD, particularly with DTI diffusivity and NODDI ISOVF measures. In contrast, NODDI ISOVF and ODI were marginally more sensitive to Aβ burden than DTI measures, not only in limbic tracts like the fornix, but the largest effects were found in the posterior internal capsule. While lower DTI FA and increased diffusivity may indicate nonspecific WM deterioration, NODDI measures can provide more insight. Lower ICVF and ODI, and higher ISOVF suggest these effects may be driven by factors such as neuronal loss, reduced tissue complexity, and greater free water which may indicate inflammation/edema, respectively.

Overall, we found a greater number of sex differences in amyloid and tau pathological markers compared to metrics of cognitive impairment, suggesting that sex-related biological factors may influence disease pathology before detectable cognitive decline. Interestingly, males showed greater disease effects than females, and not *vice versa*. It is possible that the complex interplay of genetic, hormonal (e.g., estrogen decline during menopause), and immunological factors that influence disease risk and progression in females may increase inter-individual variability and obscure group-level effects. Disentangling these overlapping influences is critical for accurately characterizing sex-specific disease trajectories and for informing the development of targeted diagnostic and therapeutic strategies.

As more ADNI4 multishell dMRI data becomes available, future work will compare multishell models that do not parse the signal, such as diffusion kurtosis imaging (DKI) and mean apparent propagator (MAP) MRI [28], to DTI and NODDI to evaluate their sensitivity in detecting AD-related WM changes. We also plan to incorporate along-tract WM analyses to refine spatial localization of observed effects. We also aim to investigate potential confounding factors of sex-specific group-level effects to unpack the complex sex-related patterns observed in AD. Integration of richer WM features across models could enhance detection of cognitive impairment, Aβ burden, and tau pathology in AD research.

## Acknowledgments

This work was supported by AARG-23-1149996, R01AG058854, RF1AG057892, R01AG087513, U01AG068057, U19AG024904, RF1NS136995, S10OD032285, U19AG024904.

